# Model-driven approach for the production of butyrate from CO_2_/H_2_ by a novel co-culture of *C. autoethanogenum* and *C. beijerinckii*

**DOI:** 10.1101/2022.10.12.511882

**Authors:** Sara Benito-Vaquerizo, Niels Nouse, Peter J. Schaap, Jeroen Hugenholtz, Stanley Brul, Ana Lopez-Contreras, Vitor A. P. Martins dos Santos, Maria Suarez-Diez

**Author notes:** These authors contributed equally to this work (Sara Benito-Vaquerizo, Niels Nouse). Correspondence: Stippeneng 4, 6708 WE, Wageningen, The Netherlands.

## Abstract

One-carbon (C1) compounds are promising feedstocks for sustainable production of commodity chemicals. CO_2_ is a particularly advantageous C1-feedstock since it is an unwanted industrial off-gas that can be converted to valuable products while reducing its atmospheric levels. Acetogens are known microorganisms that can grow on CO_2_/H_2_ and syngas converting these substrates into ethanol and acetate. Co-cultivation of acetogens with microbes that can further process such products can expand the variety of products to, for example, medium chain fatty acids (MCFA) and longer chain alcohols. Solventogens are microorganisms known to produce MCFA and alcohols via the acetone, butanol and ethanol (ABE) fermentation in which acetate is a key metabolite. Thus, co-cultivation of a solventogen and acetogen in a consortium provides a potential platform to produce valuable chemicals from CO_2_.

In this study, metabolic modelling was implemented to design a new co-culture of an acetogen and a solventogen to produce butyrate from CO_2_/H_2_ mixtures. The model-driven approach suggested the ability of the studied solventogenic species to grow on lactate/glycerol with acetate as co-substrate. This ability was confirmed experimentally by cultivation of *Clostridium beijerinckii* on these substrates in serum bottles and subsequently in pH-controlled bioreactors.

Community modelling also suggested that a novel microbial consortium consisting of the acetogen *Clostridium autoethanogenum,* and the solventogen *C. beijerinckii* would be feasible and stable. On the basis of this prediction, a co-culture was experimentally established. *C. autoethanogenum* grew on CO_2_/H_2_ producing acetate and traces of ethanol. Acetate was in turn, consumed by *C. beijerinckii* together with lactate, producing butyrate. These results show that community modelling of metabolism is a valuable tool to guide the design of microbial consortia for the tailored production of chemicals from renewable resources.

## 1 INTRODUCTION

The energy crisis and the effects of climate change have emphasised the need to accelerate the transition towards a circular bio-based economy (1). Current circular approaches focus on the use of low value feedstocks, such as biomass waste streams, for microbial conversions into commodity chemicals (2). Industrial waste gases from steel and thermal power plants can be directly used as microbial feedstocks (3). Lignocellulosic biomass can be converted into sugars by pretreatment and hydrolysis (4), but it can also be gasified to synthesis gas -syngas-, a mixture of CO, CO_2_ and H_2_ (5). In this regard, CO_2_ is an advantageous one-carbon feedstock (C1) since it can be obtained from natural and industrial sources, and can be converted into valuable products reducing the release of contaminant gases to the environment.

Acetogens are strict anaerobes that can grow on CO_2_/H_2_ and syngas as sole carbon source using the Wood-Lhungdahl metabolic pathway (6, 7). Acetogenic fermentation of C1 gases leads to acetate, ethanol, 2,3-butanediol or lactate, among other natural products (8, 9), whereas the fermentation of ethanol is well established and has been widely commercialised. (10, 11). However, alternative strategies are needed to overcome energy limitations of acetogens and the low solubility associated to C1 gases, which constrains the product spectrum and the product titres in the fermentation broth (12).

Solventogenic clostridia naturally produce mixes of acetone, butanol and ethanol (ABE) (13), that have applications as solvents, by fermentation of sugars. Some strains are able to reduce acetone into isopropanol resulting in mixtures of isopropanol, butanol and ethanol (IBE) (14). During fermentation of sugars to solvents, two phases can be observed: the acidogenesis and the solventogenesis. During acidogenesis, solventogens produce carboxylic acids (mainly acetate and butyrate), and CO_2_ and H_2_. Accumulation of carboxylic acids leads to a lower pH triggering the solventogenesis phase. In this step, the solventogen assimilates the carboxylic acids, and produces acetone, butanol and ethanol (15).

Cross-feeding strategies have been used to successfully establish synthetic microbial communities that lead to a wider product range (16, 17, 18, 19). Therefore, co-cultivation of an acetogen and a solventogen has the potential to overcome the drawbacks associated to acetogens by increasing the product spectrum. Charubin and Papoutsakis (20) recently established a co-culture of the solventogen *Clostridium acetobutylicum* and the acetogen *Clostridium ljungdahlii*. In this setup, glucose was fed to *C. acetobutylicum* which in turn produced butanol, ethanol, acetone, acetoine, CO_2_ and H_2_. CO_2_ and H_2_ were then fixed by *C. ljungdahlii* that also took up acetoin and acetone leading to production of isopropanol and 2,3-butanediol. Acetate is one of the most abundant products in acetogens (21), and while solventogenic strains cannot grow on acetate as sole carbon source, they are able to re-assimilate acetate and convert it into carboxylic acids such as butyrate, or solvents when glucose is used as co-substrate (22, 23, 24).

Therefore, in a co-culture of acetogens/solventogens on CO_2_/H_2_, butyrate could be produced as main product. Butyrate is a valuable product as it is used in many commercial applications, as a solvent, cosmetic, food, animal feed or as a precursor of pharmaceuticals (25, 26).

In this study, we have followed a model-driven approach to find an alternative route for production of butyrate from CO_2_ using a co-culture of two strains, one acetogen producing acetate from CO_2_/H_2_, and one solventogenic strain that co-metabolises acetate with an alternative carbon source to sugars into butyrate. To select the solventogenic strain, we systematically assessed growth on several carbon sources using the genome-scale metabolic models (GEM) of *C. acetobutylicum* and *C. beijerinckii*. The most promising carbon sources were experimentally tested and validated. On the basis thereof, we constructed a community model of *C. autoethanogenum* and *C. beijerinckii*, and qualitatively assessed the fermentation of CO_2_/H_2_ and the new carbon source through scenario simulations. Model predictions guided the experimental work and lead to successful establishment of this new co-culture.

## 2 MATERIALS AND METHODS

### 2.1 GEM availability and curation

The genome-scale metabolic model of *C. autoethanogenum* DSM 10061, iCLAU786 (27) was downloaded in sbml and table format and used without modification.

The genome-scale metabolic model of *C. acetobutylicum* ATCC 824, iCac802 (28) was downloaded in sbml and table format, and modifies as follows. Two reactions were defined as reversible: ATP:3-phospho-d-glycerate 1-phosphotransferase (with model identifier R0239) and Hydrogenase (R1563). Formate dehydrogenase (R1562) was removed since it was not found in the genome of *C. acetobutylicum*. Three new reactions were added in the model: Pyruvate transport (pyrt), Pyruvate exchange (EX_PYR_e), and glycerol kinase (R0426), the latter reaction was found to be present in the genome of *C. acetobutylicum* (EC 2.7.1.30; locus tag: CAC1321). Finally, we replaced the ethanol transport reaction (R1708) expressed as a proton (H^+^) symport reaction by a diffusion transport reaction. The modified version of the model (iCac803) is available at: https://gitlab.com/wurssb/Modelling/coculture_cacb.

The genome-scale metabolic model of *C. beijerinckii* NCIMB 8052, iCM925 (29) was downloaded in sbml and table format and modified as follows. Ferredoxin-NAD+ reductase (FDXNRx) and ferredoxin-NADP+ reductase (FDXNRy) reactions were removed, and Na^+^-translocating ferredoxin:NAD^+^oxidoreductase (Rnf) complex and electron-bifurcating, ferredoxin-dependent transhydrogenase (Nfn) complex, were added in the model. Transport of hydrogen reaction (Habc) was replaced by the ATPase reaction (ATPase). The EC number and genes of (S)-3-Hydroxybutanoyl-CoA:NADP+ oxidoreductase reaction (HACD1y) were modified and the EC numbers of butyryl-CoA dehydrogenase (ACOAD1) and 3-hydroxybutyryl-CoA dehydrogenase (HACD1x) as well. Finally, the reduced ferredoxin:dinitrogen oxidoreductase (ATP-hydrolysing) reaction (DNOR) was stoichiometrically balanced. The updated version of the model (iCM943) is available in the git repository.

### 2.2 *In-silico* carbon source screening in *C. acetobutylicum* and *C. beijerinckii*

We systematically assessed growth capabilities on the updated versions of the genome-scale metabolic model of *C. acetobutylicum* and *C. beijerinckii* on a wide range of carbon sources. Model simulations were done using COBRApy, version 0.15.4 (30), and Python 3.9. Growth capabilities were assessed using Flux Balance Analysis (FBA). The biomass synthesis reaction (termed ‘Biomass’ or ‘biomass’ in the respective *C. acetobutylicum* and *C. beijerinckii* models), was defined as the objective function for maximisation. Growth was considered when the growth rate was larger than 0.0001 h^-1^. For each carbon source assessed, the maximum uptake (corresponding to minus the lower bound of the associated exchange reaction denoted ‘EX xx’) was constrained to 20 mmol g_DW_^−1^ h^−1^ and the minimum uptake (corresponding to minus the upper bound of said exchange reaction) was constrained to 0.1 mmol g_DW_^−1^ h^−1^. In addition, uptake of small metabolites and ions was allowed by setting the lower bound of the corresponding exchange reaction to −1000 as described in the corresponding script in the git repository.

### 2.3 Co-culture GEM reconstruction

A compartmentalised co-culture model of *C. autoethanogenum* and *C. beijerinckii* was obtained by combining single species models iCLAU786 (27) and iCM943 (29), following a previous approach (31). In this approach, each species is considered a single compartment. The compartment associated with *C. autoethanogenum* is defined as ‘cytosol_ca’ and the compartment associated with *C. beijerinckii* is defined as ‘cytosol’. Intracellular metabolites were assigned to their respective compartment and the flag ‘_ca’ was added to the id of metabolites belonging to ‘cytosol_ca’ to distinguish them from the *C. beijerinckii* metabolites. In addition, the combined model included an extracellular compartment defined as ‘extracellular’ common to both species. Metabolites in this compartment are either secreted, metabolised or exchanged by both species, and separated from metabolites present in the cellular compartments by adding the ‘_e’ flag to the identifier. All extracellular metabolites follow the same naming system (namespace) for both species. Therefore, the same namespace was applied to metabolites secreted by both species. All metabolites present in both intracellular compartments and the extracellular compartment can be exchanged between species if the directionality favours it. Interchanged metabolites are assumed to be first transported into the extracellular compartment, before taken up by the other species using the corresponding exchange reaction. The co-culture GEM contains one biomass synthesis reaction per species, termed ‘biomass auto’ and ‘biomass_beije’ for *C. autoethanogenum* and *C. beijerinckii*, respectively. Additionally, the model includes a community biomass synthesis reaction (‘Community biomass’), which incorporates the biomass of *C. autoethanogenum* and *C. beijerinckii* in the form of metabolite ‘biomass ca’ and metabolite ‘biomass’, respectively. The combined model incorporates a transport reaction of butyrate (‘BUTex_au’) from the extracellular compartment to the intracellular compartment of *C. autoethanogenum*, and the reaction to produce butyraldehyde from butyrate (‘buttobuta’) in *C. autoethanogenum*. In addition, we have incorporated the transport reaction of acetone (‘ACETONE ca’) from the extracellular compartment to the intracellular compartment of *C. autoethanogenum*; an alcohol dehydrogenase to convert acetone to isopropanol (‘ISOBIO’); a transport reaction of isopropanol from the intracellular compartment of *C. autoethanogenum* to the extracellular compartment (‘ISOPRO ca’), and an exchange reaction of isopropanol (‘EX_IPRO_e’).

The final three-compartment co-culture model was translated into SBML level 3 version 1 (see git repository).

### 2.4 Co-culture modelling framework

Co-culture model simulations were carried out using a previously described modelling framework (31), similar to SteadyCom (32) and based on Community FBA (cFBA) (33). Specific fluxes (mmol g_DW_^−1^h^−1^) were substituted by environmental fluxes (mmol l^−1^ h^−1^), and thus, the biomass synthesis reaction of each species was changed accordingly accounting for the growth rate and biomass of each species (g_DW_ l^−1^h^−1^). The relative contribution of each species to the community biomass is calculated from the total biomass of the community and the species ratio.

### 2.5 Co-culture model simulations

In this study, we simulated hypothetical scenarios varying biomass species ratio, growth rate and substrates environmental fluxes to explore the feasible solution space of the co-culture. We selected a community biomass of 0.22 g_DW_ l^−1^ based on the average value measured for a similar co-culture of *C. autoethanogenum* and *C. kluyveri* on syngas (18). The biomass of each species was calculated based on the indicated species ratio and the community biomass. The biomass of each species was multiplied by the indicated growth rate, and the value was used to constrain the flux through the biomass synthesis reaction of each species. We assessed conditions with *C. autoethanogenum* - *C. beijerinckii* ratios ranging from 0.1-0.9 to 0.9-0.1, and growth rates from 0.005 to 0.1 h^−1^. For this exploratory analysis, we assumed equal growth rate of each species and steady-state. For each condition, we fixed the uptake rate of CO_2_ and H_2_ to 5 mmol l^−1^ h^−1^ or to 2.5 mmol l^−1^ h^−1^, covering values found in literature (18) for a similar co-culture (18). The maximum lactate uptake rate was constrained to 2.5 or 5 mmol l^−1^ h^−1^ and a minimum uptake rate of 0.1 mmol l^−1^ h^−1^ was imposed. We defined the community biomass reaction (Community biomass) as the objective function and we performed FBA to assess the feasibility of each condition. For a selected number of feasible conditions, the solution space and the set of fluxes compatible with the measured constraints was sampled using the *sample* function in the flux analysis submodule of COBRApy. Presented results are the average and standard deviation based on 5000 iterations generated at each condition (git repository).

### 2.6 Bacterial strains

The laboratory strains *Clostridium beijerinckii* NCIMB 8052 and *Clostridium acetobutylicum* ATCC 824 were stored as spore suspensions in 20% glycerol at −20 °C. Spores of *C. beijerinckii* and *C. acetobutylicum* were heat-activated for 1 min at 95 °C and 10 min at 70 °C, respectively, before inoculation. *C. autoethanogenum* DSM 10061 was kindly provided by Professor Diana Z. Sousa from the Laboratory of Microbiology, Wageningen University and Research, Wageningen, the Netherlands and was stored as vegetative cells suspended in 25% glycerol buffered with phosphate and reduced with Ti(III)citrate under anoxic conditions at −80 °C.

### 2.7 Experimental carbon source screening of *C. acetobutylicum* and *C. beijerinckii*

Initial screening experiments were performed in serum bottles containing CM2 basal medium derived from (34) with adaptations and consisting of the following components provided in final concentrations of the fully supplemented medium: 2.5 g l^−1^ yeast extract (Duchefa Biochemie), 1.0 g l^−1^ KH_2_PO_4_, 0.61 g l^−1^ K_2_HPO_4_, 1 g l^−1^ MgSO_4_·7H_2_O, 2.9 g l^−1^ ammonium acetate, 0.10 g l^−1^ 4-aminobenzoic acid, 6.6 mg l^−1^ Fe(II)SO_4_·7H_2_O, and 0.5 mg l^−1^ Na-resazurin. Acetic acid, L-lactic acid (~90%, Merck), ethanol and glycerol were added to final concentrations of 40 mM. pH was set to pH 6.1-6.2 with KOH and/or HCl. Media were made anoxic with N_2_(g) and autoclaved. D-Glucose was made anoxic and autoclaved separately and added to a final concentration of 40 mM. Media were inoculated with 4% (v/v) culture made from heat-activated spore suspension grown overnight at 37 °C in CM2 basal medium supplemented with 40 g l^−1^ d-glucose. Cultures were incubated at 37 °C and sampled at t0 and after 4 d for analysis of cell density, measured as optical density at 600 nm (OD_600_), extracellular metabolites with high-performance liquid chromatography (HPLC), and pH. Acetate co-assimilation in the initial screening experiments was determined by calculating the fraction of total carbon converted coming from acetate.

### 2.8 Cultivation experiments in pH-controlled bioreactors

pH-controlled bioreactor experiments were performed in a working volume of 2 l in Infors HT Labfors 5 bioreactors (Infors HT, Switzerland). The stirrer, set at 150 rpm, consisted of at equidistance from top to bottom a pitch-blade and two Rushton impellers. Temperature was controlled at 37 °C and pH at pH 5.5 ±0.1 using 3 M KOH and 2 M H_3_PO_4_. Foaming was controlled with Antifoam 204 (Sigma-Aldrich). In the reactors, 2.9 g l^−1^ ammonium acetate in the CM2 basal medium was replaced by 2.5 g l^−1^ ammonium sulfate. 0.75 g l^−1^ l-cysteine HCl·H_2_O was added after autoclaving.

The adapted CM2 medium was supplemented with acetic acid and L-lactic acid prior to autoclaving in pH-controlled batch fermentations of *C. beijerinckii* on different concentrations of acetate and lactate. 10 ml min^−1^ N_2_(g) was flushed across the head space to keep anoxic conditions. Reactors were inoculated with 1% (v/v) *C. beijerinckii* culture growing at 37 °C in CM2 medium supplemented with 20 g l^−1^ d-glucose and 0.75 g l^−1^ l-cysteine HCl·H_2_O.

The reactors were equipped with sinter spargers in pH-controlled fed-batch co-cultivation experiments of *C. autoethanogenum* and *C. beijerinckii* to flush 40 ml min^−1^ H_2_(g) and 10 ml min^−1^ CO_2_(g) through the medium. At t_0_ reactors were inoculated with <1% (v/v) *C. autoethanogenum* culture growing at 37 °C in CM2 medium supplemented with 10 g l^−1^ d-fructose and 0.75 g l^−1^ l-cysteine HCl·H_2_O. After establishment of growth and acetate production by *C. autoethanogenum*, reactors were inoculated with <1% (v/v) *C. beijerinckii* culture growing at 37 °C in CM2 medium supplemented with 20 g l^−1^ d-glucose and 0.75 g l^−1^ l-cysteine HCl·H_2_O. Furthermore, the continuous l-lactic acid feed was started.

Reactors were sampled at regular time intervals for analysis of cell density, extracellular metabolites with HPLC, and morphology with phase-contrast microscopy.

In pH-controlled batch fermentations with *C. beijerinckii* the overall stoichiometry was calculated by scaling the difference of the concentrations of main extracellular metabolites between t_0_ and t_end_ to the difference in lactate concentration.

In co-cultivation experiments, theoretical acetate production from CO_2_ was calculated as follows for each time point after the start of the l-lactic acid feed: The amount of lactate converted was calculated from the difference in the amount of lactate fed and calculated amount of lactate remaining in the reactor. The conversion of lactate via pyruvate yields the intermediate metabolite acetyl-CoA and CO_2_ in a 1:1 ratio. The fraction of the amount of carbon from lactate available for the formation of products was subtracted from the amount of carbon present in the produced (iso)butyrate to obtain a theoretical amount of carbon coming from a difference source than lactate, i.e., from converted acetate. This theoretical amount of converted acetate was added to the calculated amount of acetate in the reactor to get a theoretical amount of acetate produced from CO_2_. Subsequently, the stoichiometry for the production of (iso)butyrate from lactate and acetate was calculated by scaling the difference of (iso)butyrate produced, theoretical acetate converted, and lactate converted between t0 and the selected time points to lactate converted.

### 2.9 Analysis of extracellular metabolites

Concentrations of acetate, acetone, butanol, (iso)butyrate, ethanol, fructose, glycerol, glucose, and lactate were analysed with HPLC. Supernatant was mixed with an equal volume of 1 M H_2_SO_4_ with 30 mM 4-methylpentanoic or 100 mM pentanoic acid as internal standard. This was filtered through a 0.2 *μ*m regenerated cellulose filter followed by analysis on a Waters HPLC system with a Shodex KC-811 column at 65 °C, 1 ml/min 3 mM H_2_SO_4_ mobile phase, and a refractive index and UV detector.

## 3 RESULTS

Fig. 1 shows the steps for the model-driven approach followed to establish a novel co-culture of *C. autoethanogenum* and *C. beijerinckii* for the production of butyrate from CO_2_/H_2_. GEMs of solventogens were used to evaluate candidate species and possible carbon sources. After the experimental validation of the model predictions, the co-culture was successfully established.

**Figure 1.**
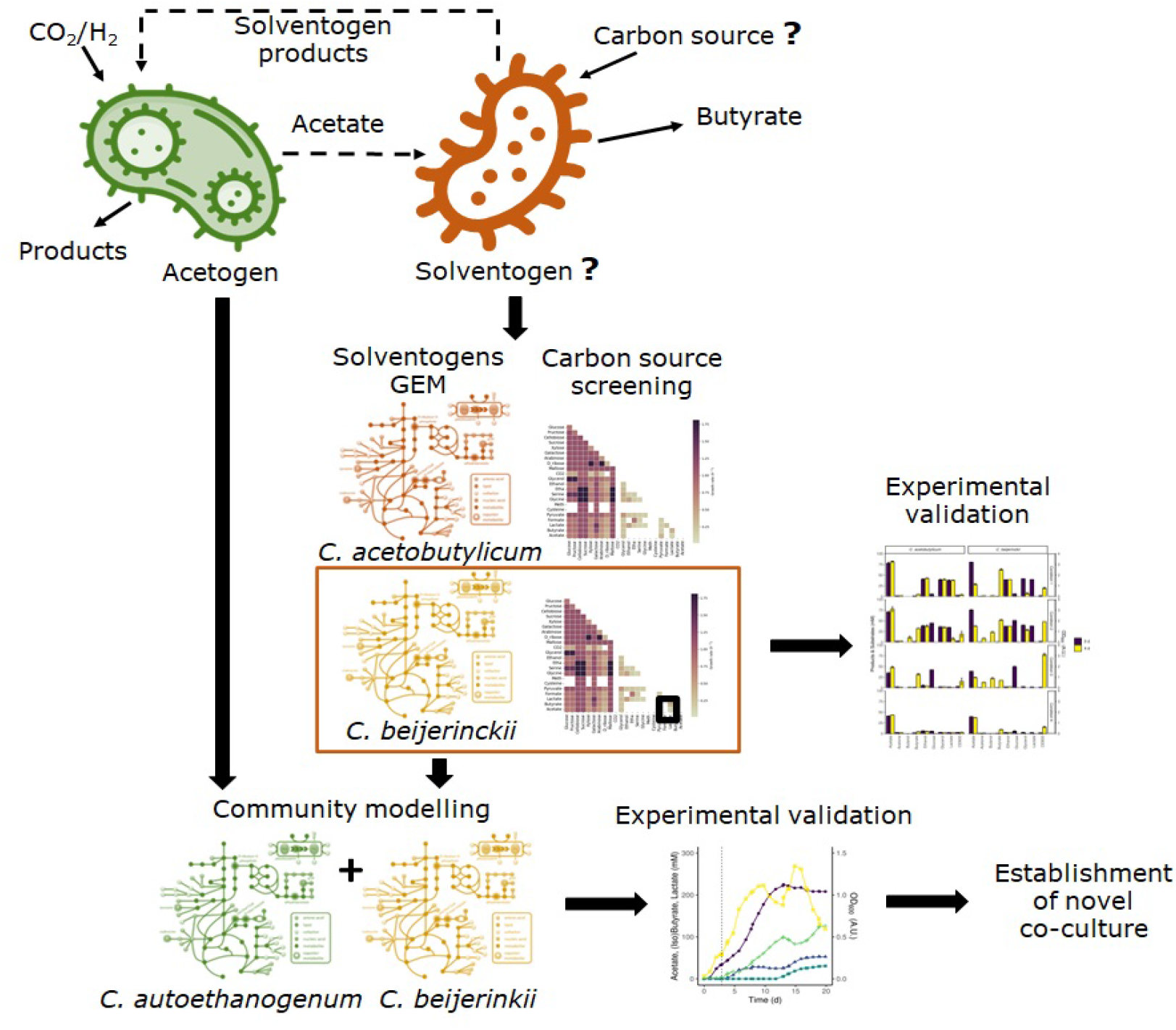
Overview of the followed methodology to establish a co-culture of an acetogen and a solventogen to produce butyrate from CO_2_/H_2_. We used genome-scale metabolic models (GEM) of two solventogens to assess growth on several carbon sources and to find the alternative carbon source that sustained growth on the solventogen with acetate. The predicted carbon sources were experimentally validated and growth was confirmed in one of the solventogens. After that, we assessed the feasibility of the co-culture of the acetogen *C. autoethanogenum* and the selected solventogen using community modelling, and finally, the co-culture was experimentally established.

### 3.1 Updated GEM of *C. autoethanogenum, C. acetobutylicum* and *C. beijerinckii*

The GEM of *C. autoethanogenum*, iCLAU786 was used as the original version (27) with no modification. The updated version of the GEM of *C. acetobutylicum*, iCac803 (28), had 1254 reactions, 1465 reactions and 803 genes, and the GEM of *C. beijerinckii*, iCM943 (29), had 881 metabolites, 941 reactions and 943 genes.

Regarding the GEM of *C. beijerinckii*, we included the Na^+^-translocating ferredoxin:NAD^+^ oxidoreductase (Rnf) complex (EC 7.2.1.2) in *C. beijerinckii*’s model. Rnf is formed by the following gene cluster: rnfC, rnfD, rnfG, rnfE, rnfA, rnfB, (locus tag: Cbei_2449, Cbei 2450, Cbei_2451, Cbei_2452, Cbei 2453 and Cbei 2454, respectively). Additionally, we have identified an electron-bifurcating, ferredoxin-dependent transhydrogenase (Nfn) complex that catalyses NADH-dependent reduced Ferredoxin:NADP+ oxidoreductase activity in *C. beijerinckii*. The Nfn complex has two subunits: NAD(P)-binding subunit, and a glutamate synthase subunit. These two subunits showed 56% to 79% identity with the Nfn subunits of *C. kluyveri, C. autoethanogenum, C. difficile* (35), and of the recently annotated *Anaerotignum neopropionicum* (36), forming two possible complexes: (Cbei_2182 and Cbei_2183) or (Cbei_0661 and Cbei_0662). To our knowledge, this is the first time the Nfn complex is reported in *C. beijerinckii* NCIMB 8052. Ferredoxin NAD+ reductase (EC 1.18.1.3) is not found in the genome of *C. beijerinckii*, and the ferredoxin NADP+ reductase (EC 1.18.1.2) showed lower % identity to the FNR of *C. acetobutylicum*, and thus, we hypothesised that this one corresponds to the NADP-binding subunit of the Nfn complex.

### 3.2 *In-silico* carbon source screening of *C. acetobutylicum* and *C.beijerinckii*

We explored growth on 25 carbon sources individually and pairs of these carbon sources with one another in *C. acetobutylicum’*s model (Supplementary data, Fig 2).

Growth was predicted on sugars, glycerol, lactate, serine and pyruvate as single carbon sources, reaching the highest growth rates on cellobiose, sucrose and maltose. As expected, acetate did not sustain growth as the sole carbon source and neither was sustained on acetone, succinate, acetoin (not shown here). Pairwise combinations of most carbon sources that led to growth as single carbon source, also led to growth in combination with an alternative carbon source. However, L-methione, L-cysteine and CO_2_ did not show growth in combination with carbon sources that sustained growth alone since the model was forced to uptake a minimum amount of each carbon source, leading in some cases, to infeasible solutions. The highest growth rates were obtained with cellobiose, sucrose or maltose in combination with serine, glycine or ethanolamine; the combination of glucose or fructose with glycerol, and xylose, or arabinose with ribose. Interestingly, acetate, in combination with lactate or glycerol, could sustain growth in *C. acetobutylicum*, as previously described for other solventogens (37, 38).

Additionally, we assessed growth on glucose, glycerol, ethanol, formate, butyrate, lactate and acetate in *C. beijerinckii*’s model (Fig. 3). As it was observed for *C. acetobutylicum, C. beijerinckii* only sustained growth on glucose, glycerol and lactate as single carbon sources. Pairwise combinations of the latter carbon sources sustained growth in combination with the rest of carbon sources. The highest growth rates were obtained with glucose in combination with glycerol or lactate. Here, acetate with lactate or glycerol also sustained growth, being the growth rate higher with addition of acetate in both scenarios.

### 3.3 Growth of *C. acetobutylicum* and *C. beijerinckii* on lactate and acetate

The substrate space provided by the model was used in an initial screening to assess growth and co-assimilation of acetate on various carbon sources by the solventogens *C. acetobutylicum* and *C. beijerinckii* (Fig. 4). On all assessed carbon sources *C. beijerinckii* grew to higher cell densities after 4 days than *C. acetobutylicum,* which is known to produce autolysins towards the end of the exponential growth phase (39). As predicted by the models (Fig. 2 and Fig. 3), neither strain grew on acetate alone (Fig. 4; Condition 4) as both strains only converted the residual metabolites from the inoculum, i.e. mainly glucose, showing the need for an additional carbon source to co-assimilate acetate. *C. acetobutylicum* could not grow on acetate with lactate, glycerol and ethanol as co-substrates, whereas the model predictions for *C. beijerinckii* were confirmed, and acetate was co-assimilated using all lactate and part of the glycerol into butyrate (Fig. 4; Condition 1). Both solventogens further reduced butyrate to butanol with the addition of glucose (Fig. 4; Condition 2). The fraction of carbon coming from acetate in the products produced by *C. beijerinckii* was not improved by the addition of glucose to the medium and was highest in the medium containing only acetate, ethanol, glycerol and lactate (Fig. 4; Condition 1-3, Table S1).

**Figure 2.**
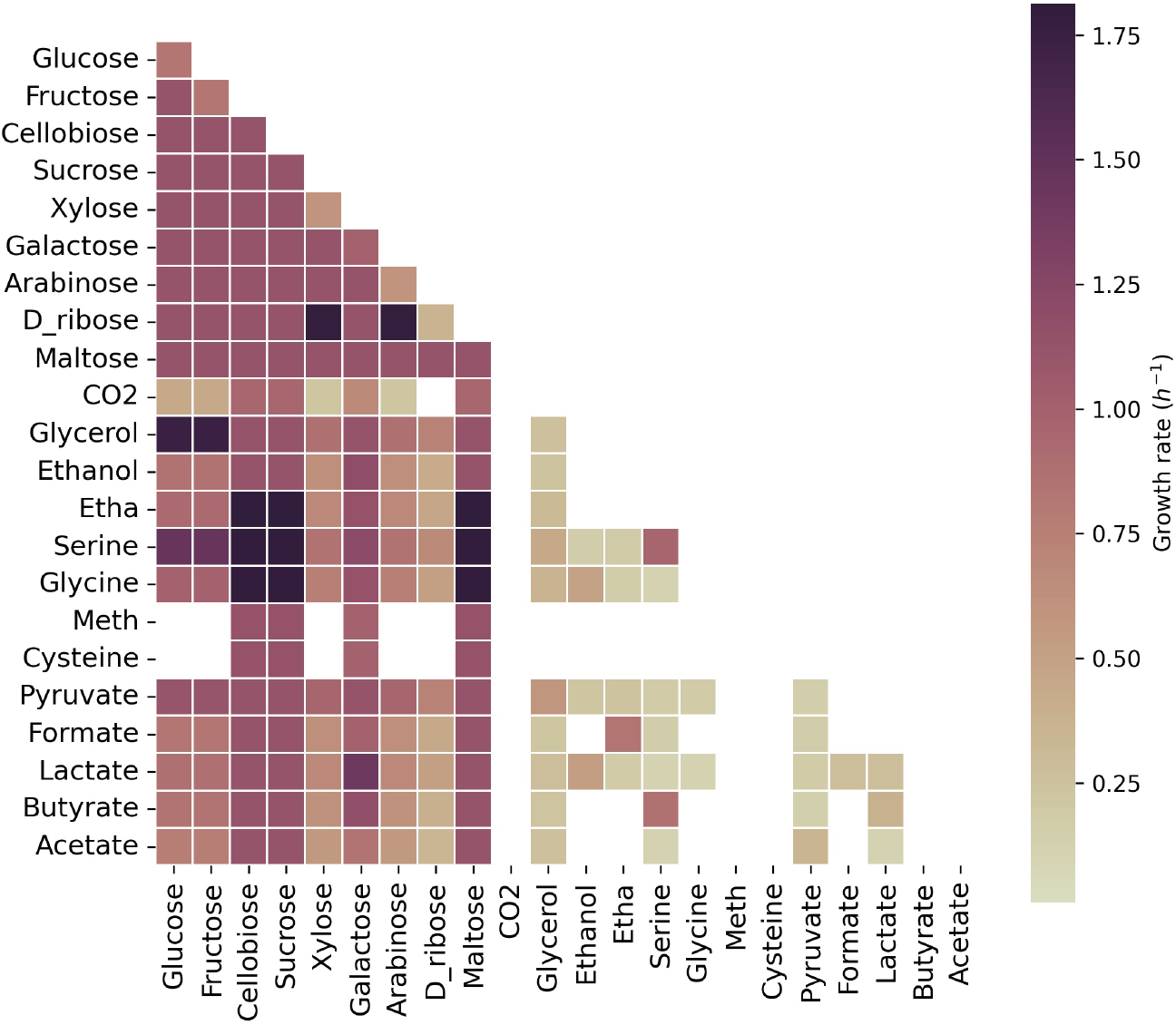
Growth capabilities predicted for *C. acetobutylicum.* The color scale indicates growth rates in h^−1^. Squares on the diagonal correspond to single carbon sources. Squares below the diagonal correspond to the combination of the carbon source indicated in x axes with the carbon source indicated in y axes. Non-colored squares means that there was no growth predicted for the specified carbon source(s). Meth: Methionine; Etha: Ethanolamine. The maximum uptake rate for each carbon source is set to 20 mmol g_DW_^−1^ h^−1^ and the minimun to 0.1 mmol g_DW_^−1^ h^−1^

**Figure 3.**
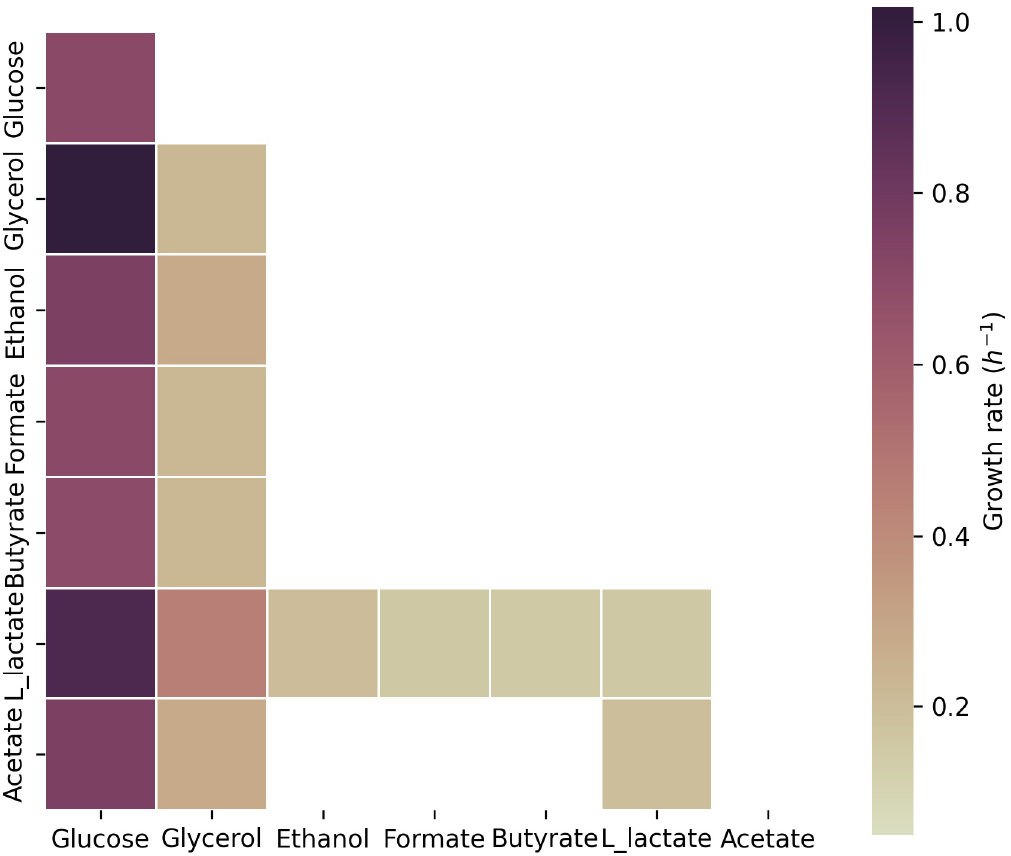
Growth capabilities predicted for *C. beijerinckii*. The color scale indicates growth rates in h^−1^. Squares on the diagonal correspond to single carbon sources. Squares below the diagonal correspond to the combination of the carbon source indicated in x axes with the carbon source indicated in y axes. Non-colored squares means that there was no growth predicted for the specified carbon source(s). The maximum uptake rate for each carbon source is set to 20 mmol g_DW_^−1^ h^−1^ and the minimun to 0.1 mmol g_DW_
^−1^ h^−1^

**Figure 4.**
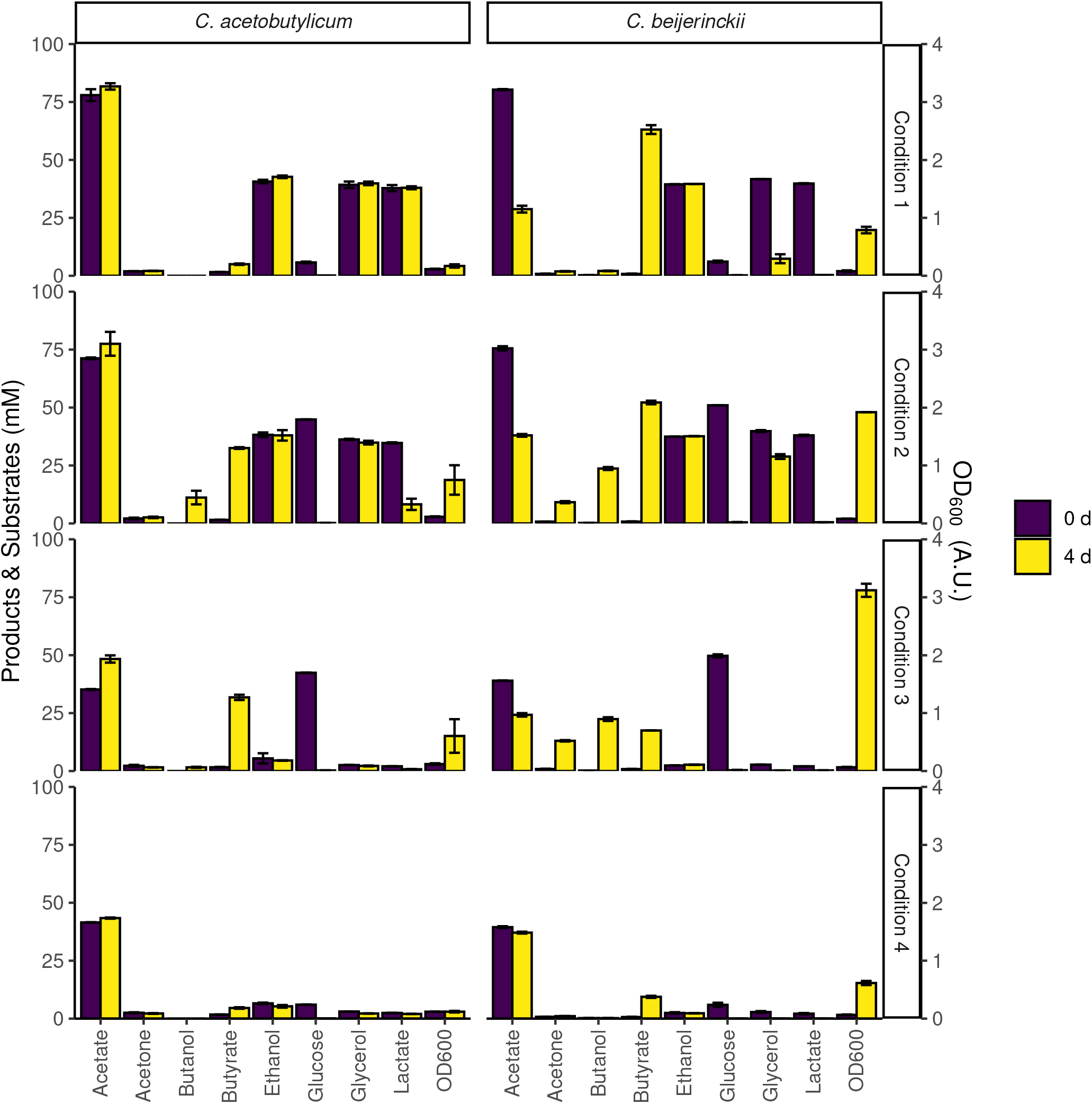
Initial screening of *C. beijerinckii* and *C. acetobutylicum* for growth on various combinations of carbon sources. The bars indicate substrate and product concentrations, and cell density at the time of inoculation and after 4 days. **Condition 1**: CM2 medium + acetic acid, ethanol, glycerol, and L-lactic acid; **Condition 2**: CM2 medium + acetic acid, ethanol, glycerol, L-lactic acid and glucose; **Condition 3**: CM2 medium + glucose. **Condition 4**: CM2 medium. Note: CM2 medium contained 38 mM acetate. Error bars show standard deviations between two cultures inoculated with the same inoculum.

Growth of *C. beijerinckii* on lactate and acetate was further explored in a bioreactor at a controlled pH of 5.5 (Fig. S1). This pH is close to the optimal pH of *C. autoethanogenum* (40), and acid re-assimilation and ABE production in *C. beijerinckii* (41, 22). Butyrate was the most abundant product with a stoichiometry of 0.6-0.7 mol of butyrate produced and 0.4-0.5 mol acetate consumed per mol of lactate consumed, similar to the stoichiometry reported for *C. saccharobutylicum* NCP262 in (37).

The screening of *C. acetobutylicum* and *C. beijerinckii* on various substrates suggested the potential of *C. beijerinckii* as solventogen, and lactate as carbon source for the re-assimilation of acetate in co-cultivation with *C. autoethanogenum*.

### 3.4 Fermentation of lactate and acetate by *C. beijerinckii*

In *C. beijerinckii*, lactate is oxidised to pyruvate via NAD-dependent L-lactate dehydrogenase (EC 1.1.1.27) encoded by the following isoenzymes: Cbei_4072, Cbei_4903 or Cbei_2789 (Fig. 5). Pyruvate is decarboxylated to acetyl-CoA via pyruvate:ferredoxin oxidoreductase (PFOR; EC 1.2.7.10, encoded by Cbei_1853, Cbei_4318 or Cbei_1458), generating reduced ferredoxin and CO_2_. Model predictions showed that reduced ferredoxin is partly spent to produce H_2_ and oxidised ferredoxin via ferredoxin hydrogenase (EC 1.12.7.2, Cbei_0327, Cbei 4000 or Cbei_3796), and partly spent to regenerate oxidised ferredoxin and NADH via the Rnf complex (EC 7.2.1.2, Cbei_2449-55), translocating Na^+^/H^+^ (42). The Rnf complex is coupled to an ATPase (EC 7.1.2.2) encoded by the cluster Cbei_0412 to Cbei_0419, that pumps in Na^+^/H^+^for energy generation.

**Figure 5.**
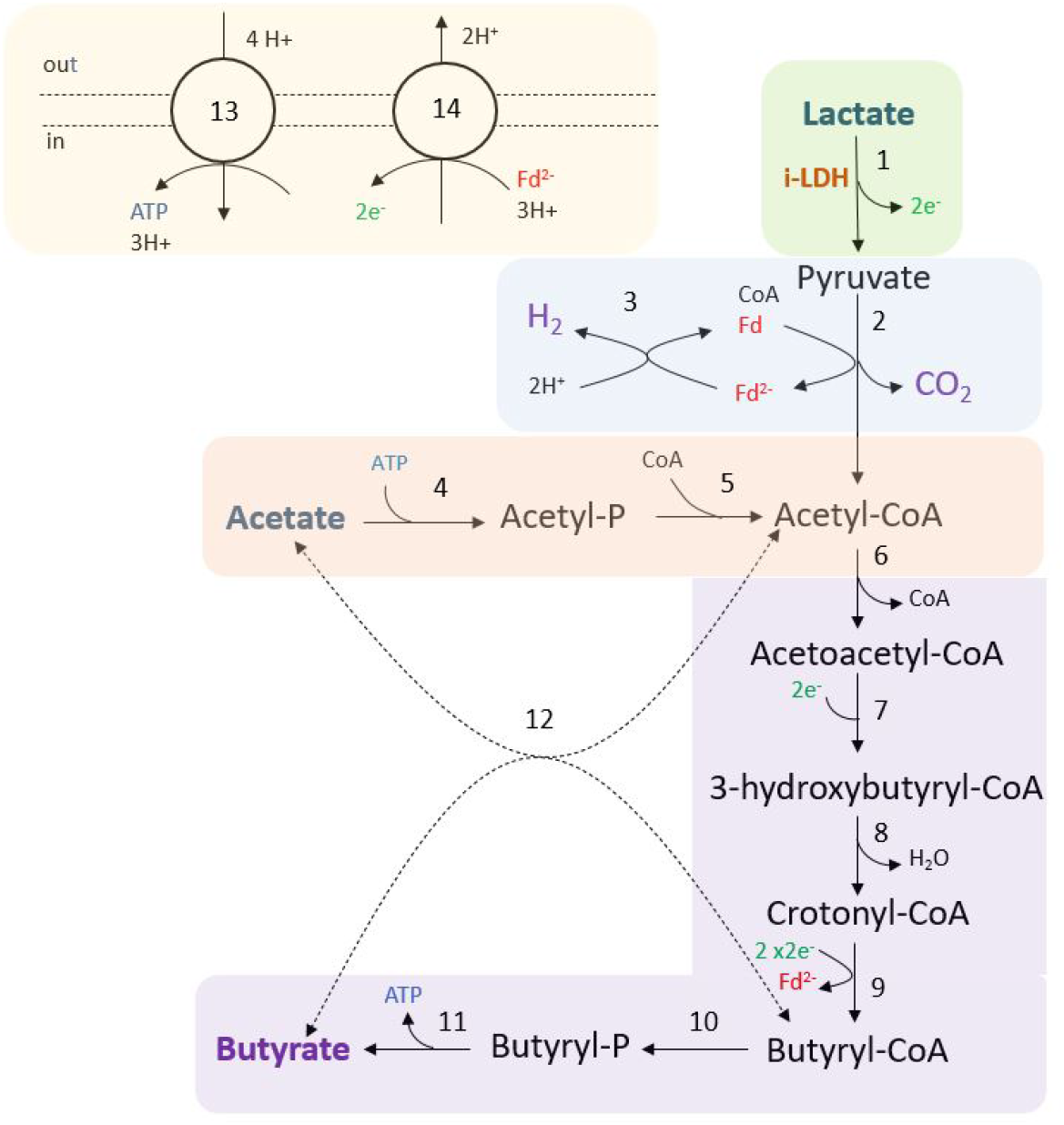
Fermentation of lactate and acetate by *C. beijerinckii*. Coloured areas designate the following modules: lactate oxidation (green); H_2_, CO_2_, and acetyl-CoA production (blue); acetate consumption (red); butyrate production (purple); and redox cofactor regeneration, and ATPase (yellow). Numbers in reactions correspond to the following enzymes and reaction ids in the model: 1, NAD-dependent L-lactate dehydrogenase (LDH_L); 2, pyruvate:ferredoxin oxidoreductase (POR4); 3, Ferredoxin hydrogenase (FDXNH); 4, acetate kinase (ACK); 5, phosphate acetyltransferase (PTA); 6, Acetoacetyl-CoA thiolase (ACACT1); 7, (S)-3-Hydroxybutanoyl-CoA:NAD+ oxidoreductase or NAD(P)-dependent acetoacetyl-CoA reductase (HACD1x or HACD1y); 8, 3-hydroxybutyryl-CoA dehydratase (3HBCD); 9, Butyryl-CoA dehydrogenase/electron-transferring flavoprotein complex (Bcd-EtfAB) (ACOAD1); 10, Butanoyl-CoA:phosphate butanoyltransferase (BCOPBT);11, Butyrate kinase (BUTK); 12, Butyryl-CoA-acetoacetate CoA-transferase (COAT2); 13, ATPase (ATPase); 14, Na^+^-translocating ferredoxin:NAD^+^ oxidoreductase complex (Rnf). Dashed lines (reaction 12) indicate that the reaction might not be the main pathway.

Model predictions suggested that acetate was consumed to acetyl phosphate (acetyl-P) investing ATP by acetate kinase (EC 2.7.2.1, Cbei_1165), and acetyl-P is converted to acetyl-CoA via phosphate acetyltransferase (EC 2.3.1.8, Cbei_3402 or Cbei_1164). As it was previously mentioned, *C. beijerinckii* produces butyrate via acetyl-CoA (43). First, two acetyl-CoA molecules are converted to one acetoacetyl-CoA by acetoacetyl-CoA thiolase (EC 2.3.1.9, Cbei_0411 or Cbei_3630). Acetoacetyl-CoA is reduced to 3-hydroxybutyryl-CoA via (S)-3-Hydroxybutanoyl-CoA:NAD+ oxidoreductase (EC 1.1.1.157, Cbei_0325) or via NAD(P)-dependent acetoacetyl-CoA reductase (EC 1.1.1.36, Cbei_5834). Then, 3-hydroxybutyryl-CoA forms crotonyl-CoA by 3-hydroxybutyryl-CoA dehydratase (EC 4.2.1.55, Cbei_2034 or Cbei_4544). Crotonyl-CoA is reduced via the butyryl-CoA dehydrogenase/electron-transferring flavoprotein complex (Bcd-EtfAB) producing reduced ferredoxin. Two complete clusters have been identified in the genome: Cbei_0322 (Bcd), Cbei_0323 (EtfB) and Cbei_0324 (EtfA) or Cbei_2035 (Bcd), Cbei_2036 (EtfB) and Cbei_2037 (EtfA). An acyl-CoA dehydrogenase (Acd) showed a 79.4% similarity with the Bcd subunit of *C. acetobutylicum* ATCC 824. Butyrate can be produced from butyryl-CoA via two routes in *C. beijerinckii*. The first route is a linear pathway in which butyryl-CoA is first converted to butyryl phosphate via butanoyl-CoA:phosphate butanoyltransferase (Ptb; EC 2.3.1.19, Cbei_0203). Butyryl phosphate is then converted to butyrate producing ATP via butyrate kinase (Buk; EC 2.7.2.7, Cbei_0204). The second route is catalysed by a butyryl-CoA-acetoacetate CoA-transferase (EC 2.8.3.9, Cbei_2654 or Cbei 2653 or Cbei_3834 or Cbei_3833 or Cbei_4614 or Cbei_4612), where the CoA moiety of butyryl-CoA is transferred to acetate producing acetyl-CoA and butyrate. However, model predictions showed that butyrate is mostly produced generating ATP via Ptb and Buk.

### 3.5 Community model simulations of *C. autoethanogenum* and *C. beijerinckii* for the fermentation of CO_2_/H_2_ and lactate

The GEM of the co-culture consisted of 2005 metabolites, 2107 reactions and 1659 genes. Community model simulations supported the co-existence of the co-culture of *C. autoethanogenum* and *C. beijerinckii* for the fermentation of CO_2_/H_2_ and lactate in a wide range of growth rates and species ratio combinations. Fig. 6 shows the feasible solution space for multiple combinations of species ratio, growth rates, CO_2_, H_2_ and lactate feed. When the maximum uptake of lactate is 2.5 mmol l^−1^ h^−1^ (Green figures), the feasibility of the co-culture becomes more limited. In these conditions, the co-culture is only feasible at low growth rates (< 0.02 h^−1^) for all species ratios, and feasible at higher growth rates (up to 0.07 h^−1^) when *C. autoethanogenum* and *C. beijerinckii* are similarly present in the community for CO_2_/H_2_ feed ratio of 0.5. The co-culture is infeasible in all conditions when the CO_2_/H_2_ feed ratio is 2, and only feasible when *C. autoethanogenum’*s presence is low and CO_2_/H_2_ feed ratio is 1. The range of feasible solutions becomes larger when lactate feed rate is 5 mmol l^−1^ h^−1^. When *C. autoethanogenum* and *C. beijerinckii* are equally present in the community, the co-culture can be established with all growth rates explored, except when the CO_2_/H_2_ feed ratio is 2, that is only feasible up to 0.04 h^−1^). Again, only at lower growth rates (< 0.02 h^−1^), the co-culture is feasible for all tested species ratio.

**Figure 6.**
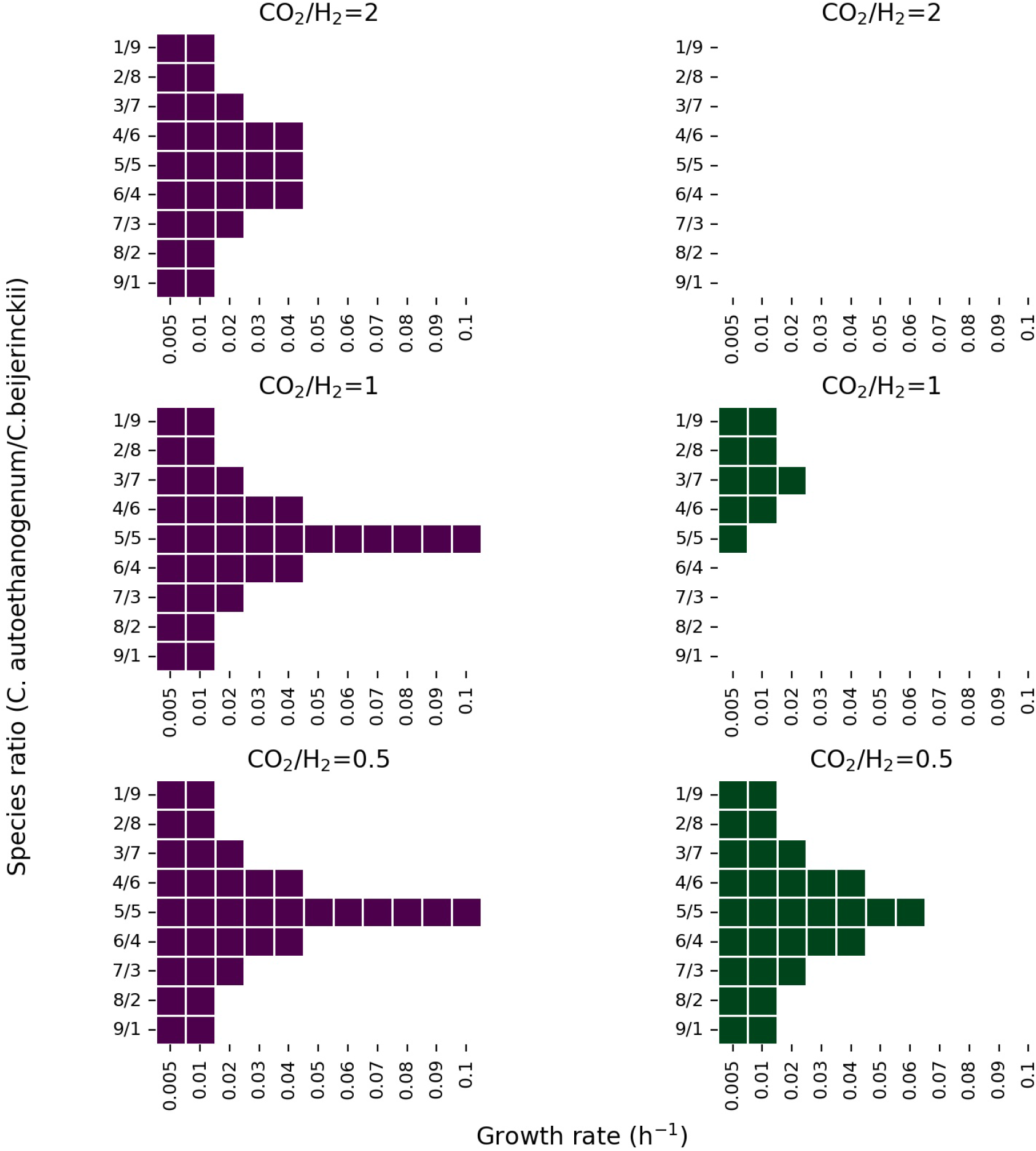
Feasible solution space of the co-culture of *C. autoethanogenum* and *C. beijerinckii* for several species ratio and growth rate combinations under different CO_2_, H_2_ and lactate feed rate. y axis shows the biomass species ratio of *C. autoethanogenumC. beijerinckii* and x axis shows growth rate in h^−1^. Colored areas indicate feasible solutions predicted by the model. Figures in purple and green show results when lactate feed rate is set to a maximum of 5 mmol l^−1^ h^−1^, and 2.5 mmol l^−1^ h^−1^, respectively. Predictions shown on the first row were obtained with a CO_2_ and H_2_ feed rate of 5, and 2.5 mmol l^−1^ h^−1^, respectively. Predictions shown on the second row were obtained with a CO_2_ and H_2_ feed rate of 5 mmol l^−1^ h^−1^, and on the third row, with a CO_2_ and H_2_ feed rate of 2.5, and 5 mmol l^−1^ h^−1^, respectively.

Fig. 7 shows the steady-state consumption and production rates observed in co-culture compared to the consumption and production rates associated to *C. autoethanogenum* or *C. beijerinckii* in the co-culture for a feasible solution indicated in the feasibility study (Fig. 6). As it can be observed, part of the acetate produced by *C. autoethanogenum* is being uptaken by *C. beijerinckii* since the steady-state production rates in co-culture are lower than the production rates by *C. autoethanogenum*. The fermentation of acetate and lactate leads to the production of butyrate in *C. beijerinckii.* A low amount of butyrate is re-assimilated by *C. autoethanogenum* and by *C. beijerinckii* and it is converted to butanol (not showing here). Furthermore, ethanol is being produced in lower amounts by *C. autoethanogenum* and *C. beijerinckii*. Model predictions also showed an exchange of CO_2_ and H_2_ from *C. beijerinckii* to *C. autoethanogenum*. *C. beijerinckii* produces CO_2_ and H_2_ that are taken up by *C. autoethanogenum*, since the flux through *C. autoethanogenum* is higher than the flux through the exchange reaction in the co-culture. Model predictions suggested that acetate consumption by *C. beijerinckii* varied depending on the lactate feed rate, being lower when the lactate feed rate was higher than 2.5 mmol l^−1^ h^−1^ (≈5 mmol l^−1^ h^−1^). A higher lactate feed rate also led to more CO_2_ and H_2_ in *C. beijerinckii*, and thus, to more gases being recirculated and consumed by *C. autoethanogenum* producing more acetate (git repository).

**Figure 7.**
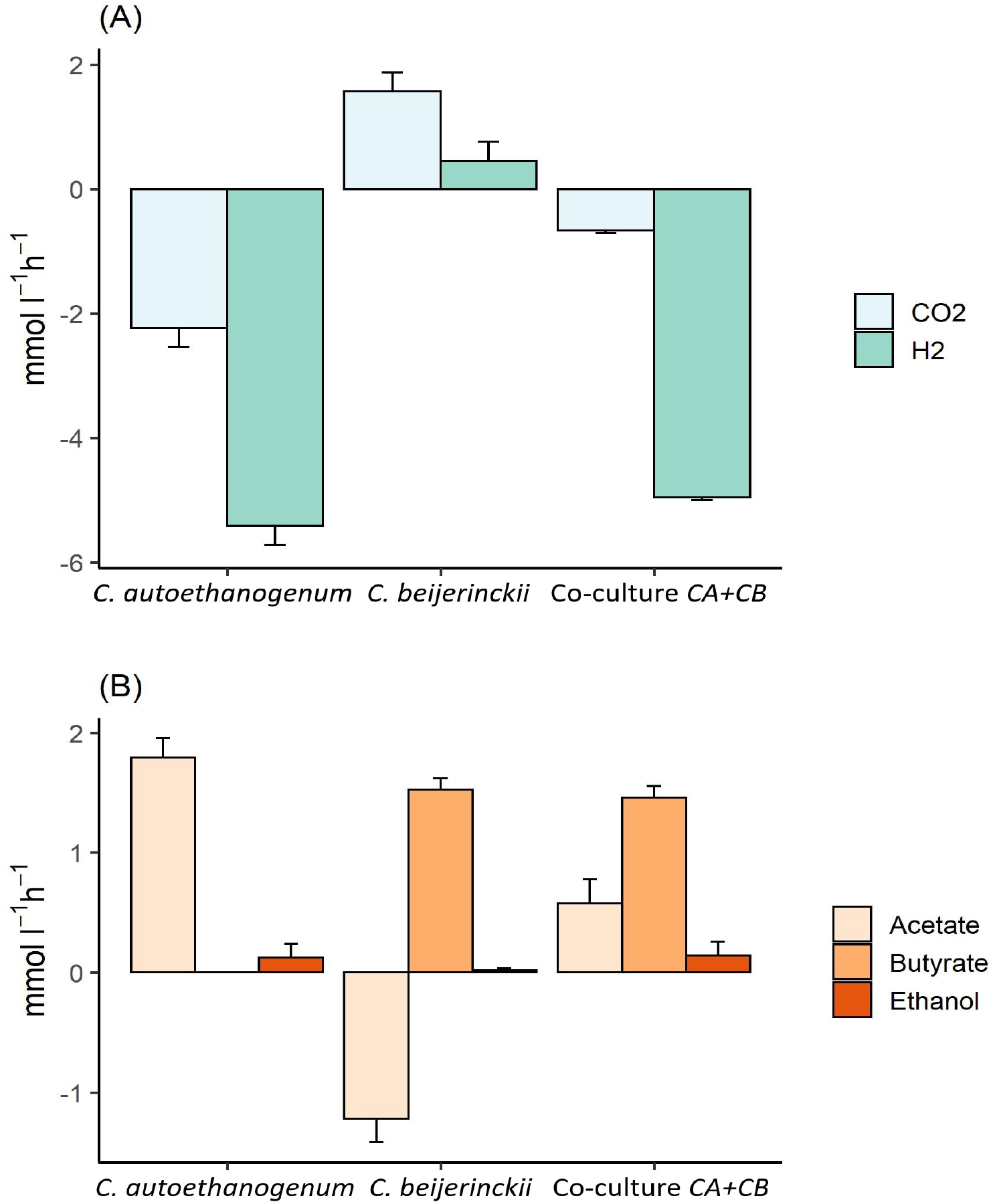
Steady-state production and consumption rates of the main substrates and products predicted by the community model of *C. autoethanogenum* and *C. beijerinckii*. X axis indicates the species associated to the illustrated fluxes, and y axis shows uptake (negative) or production (positive) fluxes in mmol l^−1^ h^−1^. (A) panel shows CO_2_ and H_2_ production or consumption rates, and (B) shows the production or uptake of acetate, butyrate and ethanol. Modelled uptake and production rates are indicated for each species, indicated by *C. autoethanogenum* and *C. beijerinckii* and for the co-culture. Growth rate is set to 0.02 h^−1^; biomass species ratio is set to 1:1; maximum and minimum lactate uptake rate is set to 2.5 and 0.1 mmol l^−1^ h^−1^, and the maximum and minimum uptake of CO_2_ and H_2_ is set to 5 and 0.5 mmol l^−1^ h^−1^.

Furthermore, we observed traces of formate, butanediol, acetone, isopropanol and butanol (see git repository).

### 3.6 Fed-batch fermentation of CO_2_/H_2_ and lactate by the novel co-culture of *C. autoethanogenum* and *C. beijerinckii*

Production of butyrate from CO_2_/H_2_ and the co-substrate lactate by the modelled co-culture of *C. autoethanogenum* and *C. beijerinckii* was experimentally verified with a pH-controlled fed-batch fermentation (Fig. 8, see also Fig S2 for additional experiments). Initially, *C. autoethanogenum* was grown solely on a continuous CO_2_/H_2_ feed and after 3 d the OD_600_ had reached a value of 0.29 and the acetate concentration a value of 34 mM. This acetate concentration was considered sufficient to support *C. beijerinckii*, therefore this strain was added and the L-lactic acid feed was started. The rate of the L-lactic acid feed was set lower than the rate of acetate production from CO_2_/H_2_ by *C. autoethanogenum* to prevent complete depletion of acetate. Upon inoculation with *C. beijerinckii* and the start of the L-lactic acid feed, a continued growth phase was observed till 10 d in which butyrate was produced up to 28 mM.

**Figure 8.**
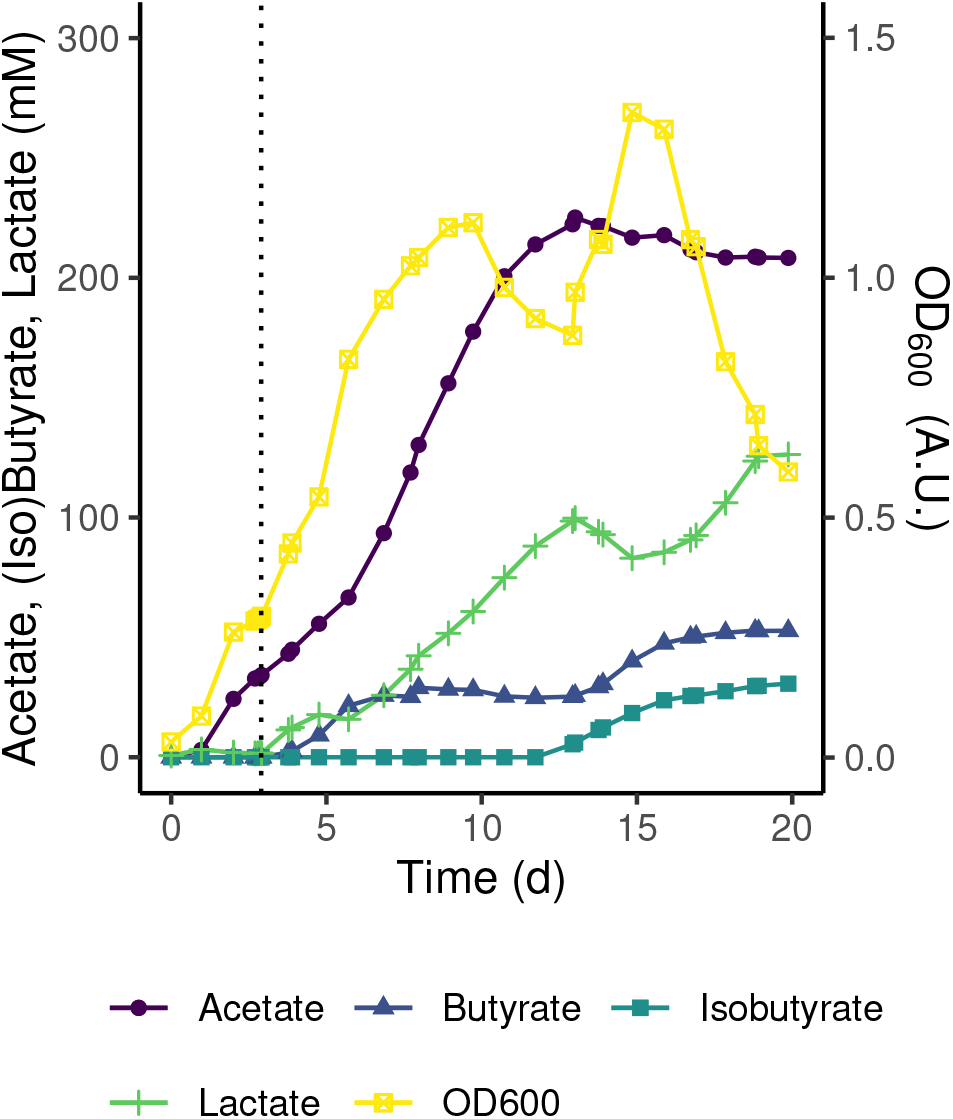
pH-controlled fed-batch fermentations of *C. autoethanogenum* - *C. beijerinckii* co-culture on 1:4 CO_2_/H_2_ with l-lactic acid feed. Concentrations of the main substrates, products and cell density are shown. At t_0_ cultures were inoculated with *C. autoethanogenum*. The dotted black line marks inoculation with *C. beijerinckii* and the start of the l-lactic acid feed. L-lactic acid was fed at a rate of 3 ml d^−1^ till 19 d. Traces of ethanol (3 d - 8 d, max. 4.6 mM at 7 d and 13d - 20 d, max. 5.6 mM at 20 d), butanol (14 d - 20 d, max. 3.1 mM at 20 d) and glucose from the *C. beijerinckii* inoculum (<1 mM at 3 d) were detected. pH was controlled at pH 5.5 ± 0.1.

A theoretical acetate production from CO_2_ was calculated from which the corresponding stoichiometry for butyrate production at each time point was calculated (Fig. S2). Between 4 d and 7 d, during butyrate production in the first growth phase, for each mol of consumed lactate 0.2-1 mol acetate was re-assimilated and 0.5-0.6 mol butyrate was produced.

The drop in cell density observed between 10 d and 13 d could be explained by the accumulation of biomass observed at the reactor wall above the fermentation medium from 6 d onward (data not shown). No production of butyrate was observed in this period. Until the end of this phase, the consortium consisted almost entirely of vegetative cells (data not shown). These cells could not be assigned to either species as the morphologies of *C. autoethanogenum* and *C. beijerinckii* could not be clearly distinguished by microscopic observation with phase-contrast microscopy.

After this adaptation period, a second growth phase was observed between 13 d and 15 d coinciding with a larger fraction of sporulating cells in the culture and the co-production of butyrate and isobutyrate to final concentrations of 53 mM and 31 mM, respectively (Fig. 8 and Fig. S2). The production of isobutyrate by the consortium was not predicted by the models and will be further investigated in a follow-up research. The calculated stoichiometry indicated a shift towards the conversion of lactate during this second growth and production phase (Fig. S2).

### 3.7 Analysis of substrate consumption and product formation by the co-culture model

*C. autoethanogenum* takes-up CO_2_ and H_2_ through the Wood-Ljungdahl pathway, where H_2_ is used as an electron donor for CO_2_ reduction to acetyl-CoA (Fig. 9). Acetyl-CoA is mainly converted to acetate producing ATP, and ethanol. In addition, traces of 2,3-butanediol, formate and lactate were predicted by the model (not showing here). Part of the acetate was in turn taken-up by *C. beijerinckii* together with the external lactate feed, following the metabolism described in section 3.3. Ethanol is produced by *C. autoethanogenum* and by *C. beijerinckii,* as observed in the experiments. Model simulations suggested the production of butyrate, CO_2_ and H_2_ by *C. beijerinckii*, and traces of acetone and butanol. We observed that most of the CO_2_ and H_2_ produced by *C. beijerinckii* was metabolised by *C. autoethanogenum* (Fig. 7). Furthermore, we observed traces of isopropanol formed from the conversion of the assimilated acetone by *C. autoethanogenum* through an alcohol dehydrogenase. The community model suggested that butanol was produced by *C. beijerinckii* and by *C. autoethanogenum* (Fig. 9). As Diender et al. already observed (17), butyrate could be exchanged between *C. beijerinckii* and *C. autoethanogenum*, and be converted to butanol by an alcohol dehydrogenase and the aldehyde ferredoxin oxidoreductase.

**Figure 9.**
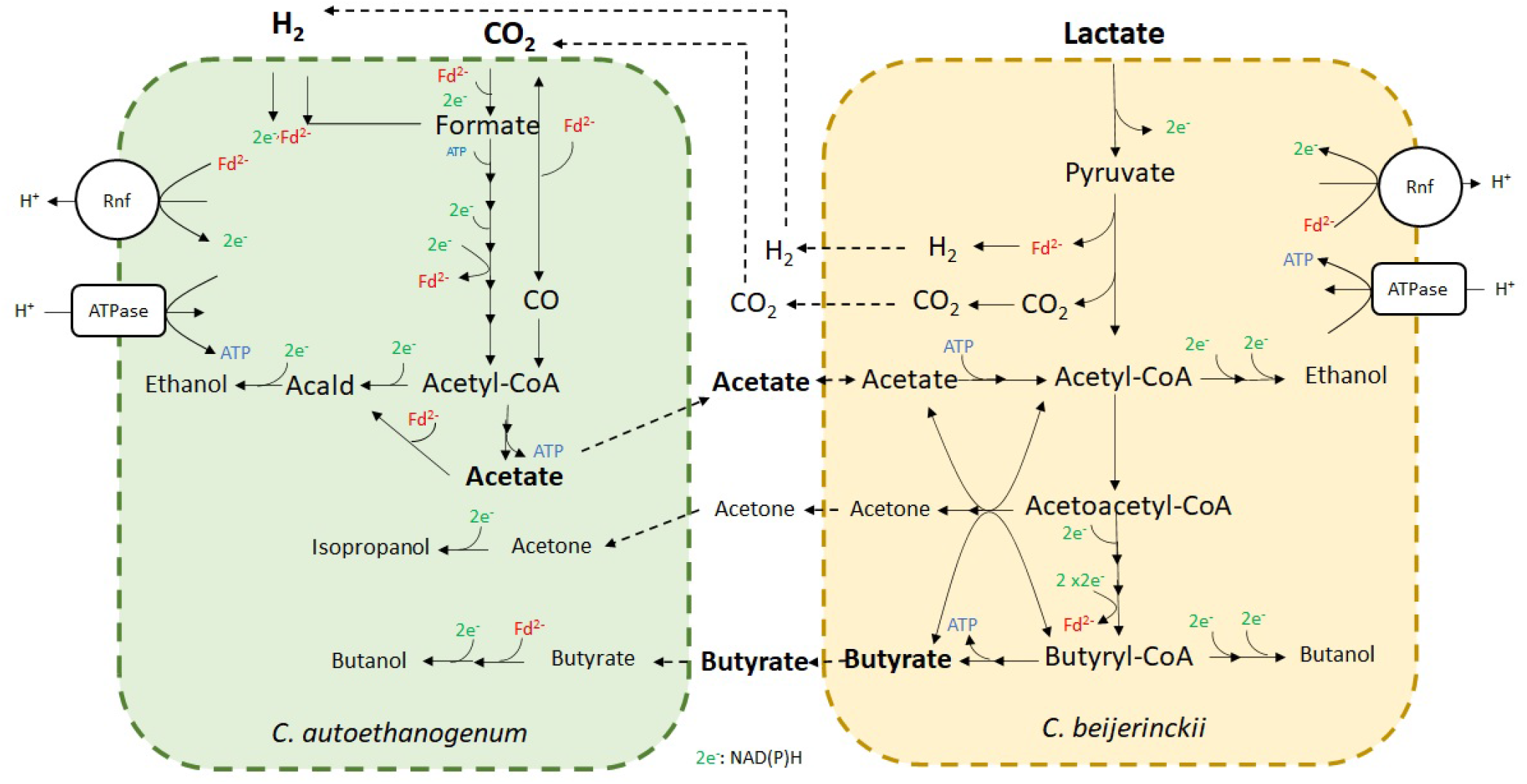
Fermentation of CO_2_/H_2_ and lactate by the novel co-culture of *C. autoethanogenum* and *C. beijerinckii*. Metabolites in bold indicate substrates and main products. Metabolites in smaller letter size indicate minor products. Arrows indicate the flux direction. Dashed lines indicate transport reactions of metabolites from extracellular compartment to intracellular compartment of the indicated microbe, and viceversa.

In addition, the model predicts traces of formate and lactate produced by *C. autoethanogenum* being assimilated by *C. beijerinckii* (not shown here).

Fig. S3 represents the metabolic profile of the novel co-culture if glucose would be the additional carbon source, instead of lactate. As observed in fig. 4, the addition of glucose would lead to an increase of solvent production (ethanol, acetone, butanol) in *C. beijerinckii*, since there are more reducing equivalents when glucose is converted to pyruvate. Acetate would still be the main product in *C. autoethanogenum* and ethanol would be produced in minor amounts. Part of acetate could be metabolised by *C. beijerinckii*, but also produced together with butyrate during the acidogenesis phase. Once the pH drops enough, the acids could be partly re-assimilated during the solventogensis phase, producing the solvents. Acetone and butyrate could be partly taken up by *C. autoethanogenum* producing isopropanol and more butanol. Possibly, part of CO_2_ and H_2_ produced by *C. beijerinckii* would be consumed by *C. autoethanogenum* as in the co-culture on lactate.

## 4 DISCUSSION

Genome-scale metabolic models are mathematical representations of the metabolism and have been successfully employed to gain insights into metabolic capabilities of single species (44, 36, 27, 28, 29), and to elucidate possible strategies to optimise microbe(s) performance in mono- and co-cultivation (45, 46, 31, 47). In this study, the use of constrained-based modelling has been key to design an alternative way to produce butyrate from CO_2_/H_2_ and lactate. We have proven the capacity of *C. beijerinckii* NCIMB 8052 strain to grow on lactate and acetate as the sole carbon and energy source. Learning from this capacity of *C. beijerinckii,* and moved by the need to upcycle sustainable feedstocks, we have established a novel synthetic co-culture of *C. autoethanogenum* and *C. beijerinckii* for the fermentation of CO_2_/H_2_ and lactate into butyrate.

The use of lactate as alternative carbon source by *C. beijerinckii* as co-substrate with acetate opens new possibilities for production of butyrate. Acetate is the most abundant product in gas fermentation and therefore, it is key in the establishment of the co-culture. Lactate has been reported to be a minor fermentation product by acetogens grown on syngas or CO_2_/H_2_ (9, 48), but a major fermentation product by acetogens grown on sugars (49). Therefore, acetogens can be engineered to increase lactate production on CO_2_/H_2_ (50), or engineered to grow on CO_2_, instead of sugars, producing lactate, and thus, produce butyrate in co-cultivation with *C. beijerinckii*. Furthermore, lactate can be obtained as side-stream in dairy industry, in almost all stored AgriFood as a result of a rotting (spontaneous fermentation) process, from storage using ensiling (51), or from the fermentation of grass (52), which could be alternative sources when establishment this co-culture.

Additionally, the use of this co-culture could increase carbon recycling and e^−^ transport, since model predictions indicated that CO_2_ and H_2_ produced by *C. beijerinckii* were almost fully re-assimilated by *C. autoethanogenum* (Fig 7), reducing carbon footprint. This is interesting to contemplate in future approaches where an organism is able to produce the H_2_ needed for the CO_2_ assimilation. Besides solventogenic clostridia, other anaerobic bacterial species have been described that produce H_2_ from sugar fermentation at high yields (53), which opens new alternatives for more efficient co-cultures without the need of external H_2_.

Model predictions showed slow growth on lactate and acetate in *C. acetobutylicum*, but this was not confirmed by experiments, where lactate was hardly consumed, and acetate was produced rather than consumed (Fig. 4; condition 1). Diez-Gonzalez et al. (37) showed growth on lactate and acetate in *Clostridium saccharobutylicum* NCP262, formerly known as *C. acetobutylicum* P262 (54). They analysed cell extracts grown on lactate and acetate and they observed both, NAD-dependent lactate dehydrogenase (d-LDH) activity, and a NAD-independent lactate dehydrogenase activity (i-LDH). d-LDH regulated the conversion of pyruvate to lactate and required fructose-1,6-biphosphate to be active (55, 37). The i-LDH regulated the conversion of lactate to pyruvate and had double the activity over d-LDH (Fig. 5). In addition, i-LDH activity decreased fourfold when glucose was added to cultures on lactate and acetate. However, this does not occur in *C. acetobutylicum*, where lactate conversion only occurs with addition of glucose (Fig. 4; Condition 2), suggesting instead, the activation of i-LDH in the presence of glucose. Interestingly, *C. beijerinckii* NCIMB 8052 showed 87.7% and *C. acetobutylicum* strain ATCC 824 showed 57% similarities with LDH of *C. saccharobutylicum* NCP262. Keis and collaborators (54) showed that *C. saccharobutylicum* NCP262 was more similar to *C. beijerinckii* NCIMB 8052 strain than to *C. acetobutylicum* ATCC 824 strain. Therefore, we hypothesised that *C. beijerinckii* would have a i-LDH activity comparable to *C. saccharobutylicum*, and *C. acetobutylicum* ATCC 824 is regulated differently.

Model predictions showed a high production of butyrate, acetate, and traces of ethanol, acetone, butanol, isopropanol, 2,3-butanediol, and formate. Fed-batch experiments also showed butyrate and acetate as major fermentations products, and ethanol and butanol as minor fermentation products. Charubin et al. (20) observed production of 2,3-butanediol from the assimilation by *C. ljungdahlii* of the acetoin produced by *C. acetobutylicum*. However, acetolactate descarboxylase was only found in *C. autoethanogenum*’s genome, and not in *C. beijerinckii*’s genome, and thus, acetoin could not be produced by the solventogen. Lactate degradation results in less NAD(P)H available, and therefore, the production of solvents is lower compared to the standard ABE fermentation on sugars (56, 20). In contrast, this co-culture has a relatively high butyrate production (up to 53 mM). Furthermore, mono-culture experiments on lactate and acetate led to higher concentrations of butyrate than the reported co-assimilation of glycerol and acetate by *C. beijerinckii* (≈ 20 mM) (38), and than the co-assimilation of lactate and acetate by *C. saccharobutylicum* (37) (≈ 20 mM).

The model built indicated that *C. beijerinckii* could grow on lactate as the sole carbon and energy source, as it was recently observed (57), but the growth rate is favoured with addition of acetate (Fig. 3), as it was previously shown (37). The addition of acetate favours lactate uptake, since acetyl-CoA pool increases with addition of acetate as co-substrate, and thus, more acetyl-CoA would be converted into butyrate producing more ATP. Co-culture fed-batch experiments showed however, accumulation of acetate in the fermentation broth, showing that not all acetate produced by *C. autoethanogenum* was consumed by *C. beijerinckii*, as indicated by the model, and possibly some acetate could also be produced by *C. beijerinckii*.

We should note that the deployed modelling approach predicts steady-state production or consumption rates, and thus, we cannot compare the results quantitatively with bioreactor data, which consist of concentrations over time. Instead our study should be seen as an exploratory study assessing the feasibility of the co-culture. Future optimization of this co-culture could integrate current experimental data into dynamic modelling approaches to gain better insights into the concentration profiles over time. These results show that community modelling of metabolism is a valuable tool to guide the design of microbial consortia for the tailored production of important chemicals from renewable resources. It thereby expands the space of options to possibly accelerate the transition to a biobased economy.

## CONFLICT OF INTEREST STATEMENT

VMds has interests in LIfeGlimmer and JH has interests in NoPalm Ingredients BV. The authors declare that they have no known competing financial interests or personal relationships that could have influenced the work reported in this paper.

## AUTHOR CONTRIBUTIONS

SBV and NN conceived and designed the study, and drafted the manuscript. SBV constructed the models and performed model simulations and data analysis. NN performed the experiments and data analysis. MSD, AL, JH, SB, PJ conceived, designed and supervised the research. VMdS acquired project funding, conceived, designed and supervised the research. All authors reviewed and edited the study. All authors read and approved the content of the submitted version.

## FUNDING

The research leading to these results has received funding from the Netherlands Science Foundation (NWO) under the Programme ‘Closed Cycles’ (Project nr. ALWGK.2016.029) and the Netherlands Ministry of Education, Culture and Science under the Gravitation Grant nr. 024.002.002.

## SUPPLEMENTAL DATA

### DATA AVAILABILITY STATEMENT

The data generated for this study can be found in the following git repositorty:

https://gitlab.com/wurssb/Modelling/coculture_cacb.

## Notes

https://gitlab.com/wurssb/Modelling/coculture_cacb

