## Supplementary Material for "Model-driven approach for the production of butyrate from CO_2_/H_2_ by a novel co-culture of *C. autoethanogenum* and *C. beijerinckii*"

### 1 SUPPLEMENTARY TABLES AND FIGURES

#### 1.1 Tables

**Table S1.** Consumption/production of metabolites and co-assimilation of acetate by *C. acetobutylicum* and *C. beijerinckii* growing on various carbon sources. **Condition 1**: CM2 medium + acetic acid, ethanol, glycerol, and L-lactic acid; **Condition 2**: CM2 medium + acetic acid, ethanol, glycerol, L-lactic acid and glucose; **Condition 3**: CM2 medium + glucose. **Condition 4**: CM2 medium. Note: CM2 medium contained 38 mM acetate. Negative and positive values show conversion and production, respectively. Acetate co-assimilation is shown as the percentage of carbon converted coming from acetate (C from acetate). NA: Not available.

| | ***C. acetobutylicum*** | | | | ***C. beijerinckii*** | | | |
|---|---|---|---|---|---|---|---|---|
| **Condition** | **1** | **2** | **3** | **4** | **1** | **2** | **3** | **4** |
| **Difference (mM)** | | | | | | | | |
| Acetate | 4 | 6 | 13 | 2 | -52 | -38 | -15 | -2 |
| Acetone | 0 | 0 | -1 | 0 | 1 | 8 | 12 | 0 |
| Butanol | 0 | 11 | 2 | 0 | 2 | 23 | 22 | 0 |
| Butyrate | 3 | 31 | 30 | 3 | 62 | 51 | 17 | 9 |
| Ethanol | 2 | 0 | -1 | -1 | 0 | 0 | 0 | 0 |
| Glucose | -6 | -45 | -42 | -6 | -6 | -51 | -49 | -6 |
| Glycerol | 1 | -1 | 0 | -1 | -34 | -11 | -3 | -3 |
| Lactate | 0 | -26 | -1 | 0 | -39 | -37 | -2 | -2 |
| C from Acetate (%) | NA | NA | NA | NA | 29 | 14 | 9 | 7 |

### 1.2 Figures

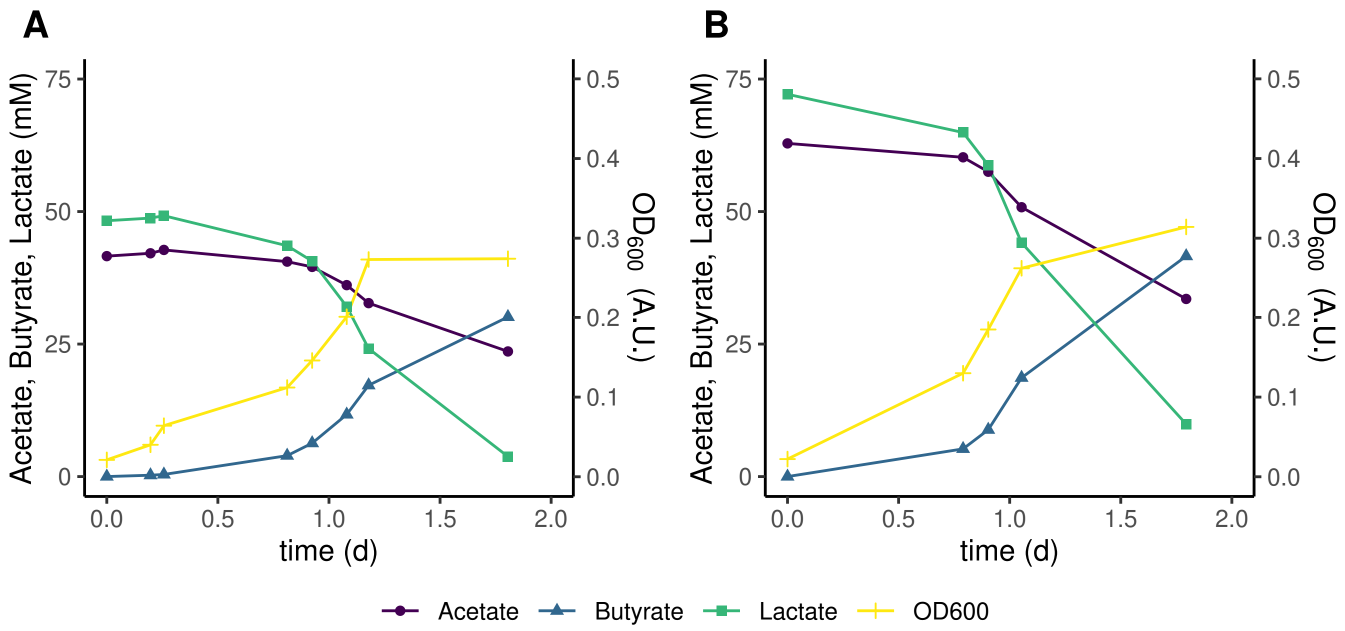


**Figure S1.** pH-controlled batch fermentations of *C. beijerinckii* growing on L-lactic acid and acetic acid at a 1:1 ratio at pH 5.5 ± 0.1. Concentrations of the main substrates, products and cell density are shown for two biologically independent replicates with different starting concentrations. Traces of ethanol and butanol (<1 mM), and glucose from the inoculum (1 mM) were detected in both conditions. Overall reaction stoichiometries for lactate, acetate and butyrate were -1, -0.40 and 0.68 for (**A**), and -1, -0.47 and 0.67 for (**B**), respectively.

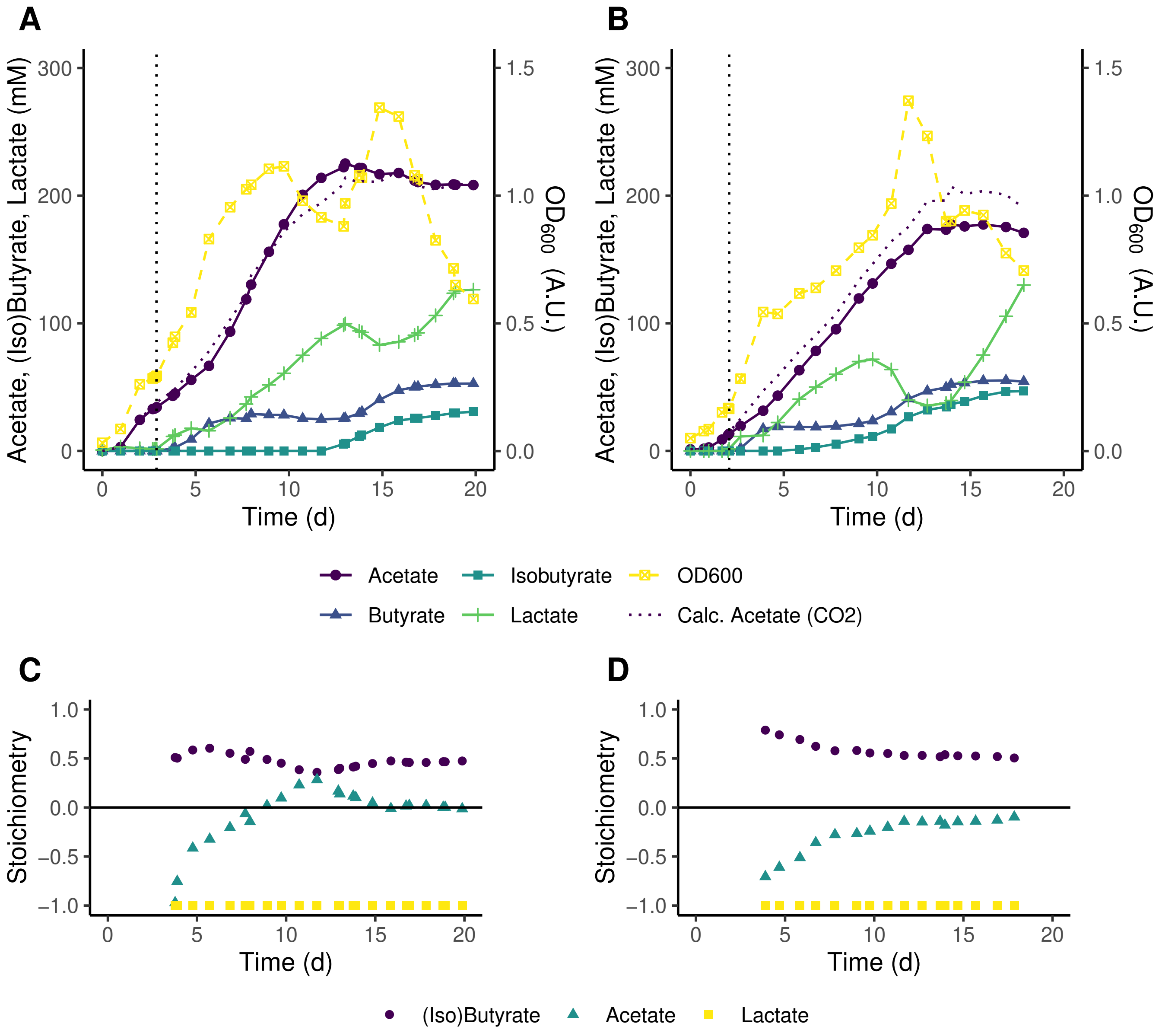


**Figure S2.** pH-controlled fed-batch fermentations of *C. autoethanogenum* - *C. beijerinckii* co-culture on 1:4 $CO_2/H_2$ with L-lactic acid feed at pH 5.5 ± 0.1. Concentrations of the main substrates, products and cell density are shown for two biologically independent replicates in (**A**) and (**B**) and corresponding stoichiometries in (**C**) and (**D**), respectively. At $t_0$ cultures were inoculated with *C. autoethanogenum*. The dotted black line in (**A**) and (**B**) marks inoculation with *C. beijerinckii* and the start of the L-lactic acid feed. In (**A**) L-lactic acid was fed at a rate of 3 ml $d^{-1}$ till 19 d and in (**B**) at a rate of 3 ml $d^{-1}$ till 14 d and 6 ml $d^{-1}$ from 14 d till 18 d. A theoretical acetate concentration produced from $CO_2$ was calculated (Calc. Acetate ($CO_2$)). In (**A**) traces of ethanol (3 d - 8 d, max. 4.6 mM at 7 d and 13d - 20 d, max. 5.6 mM at 20 d), butanol (14 d - 20 d, max. 3.1 mM at 20 d) and glucose from the *C. beijerinckii* inoculum (<1 mM at 3 d) were detected. In (**B**) traces of ethanol (1 d - 18 d, max. 5.1 mM at 15 d) and butanol (3 d - 18 d, max. 6.3 mM at 17 d) were detected.

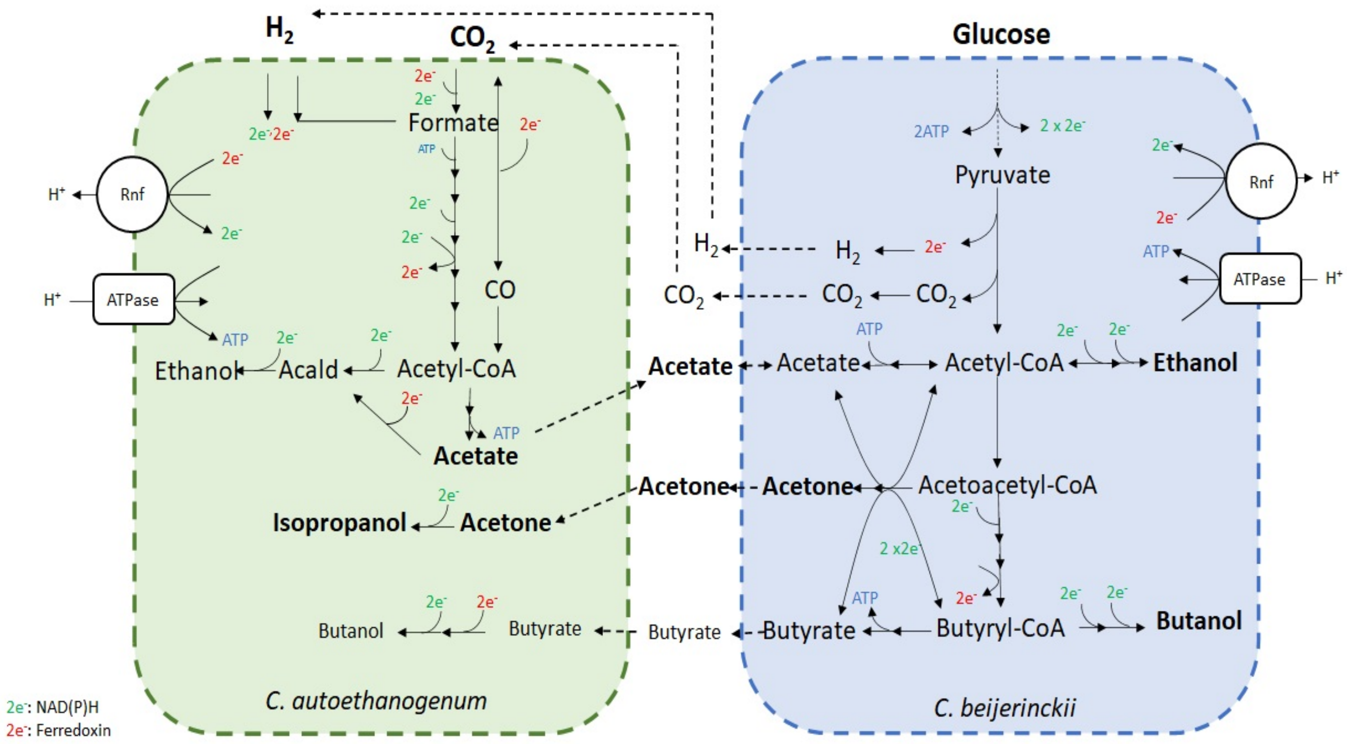


**Figure S3.** Hypothetical fermentation of $CO_2/H_2$ and glucose by *C. autoethanogenum* and *C. beijerinckii*. Metabolites in bold indicate substrates and main products. Metabolites in smaller letter size indicate minor products. Arrows indicate the flux direction. Dashed lines indicate transport reactions of metabolites from extracellular compartment to intracellular compartment of the indicated microbe, and viceversa.
